# *Listeria monocytogenes* virulence factors are secreted in biologically active Extracellular Vesicles

**DOI:** 10.1101/210906

**Authors:** Carolina Coelho, Lisa Brown, Maria Maryam, Meagan C. Burnet, Jennifer E. Kyle, Heino M. Heyman, Raghav Vij, Jasmine Ramirez, Rafael Prados-Rosales, Gregoire Lauvau, Ernesto S. Nakayasu, Nathan Ryan Brady, Anne Hamacher-Brady, Isabelle Coppens, Arturo Casadevall

## Abstract

Outer membrane vesicles produced by Gram-negative bacteria have been studied for half a century but the possibility that Gram-positive bacteria secreted extracellular vesicles (EVs) was not pursued due to the assumption that the thick peptidoglycan cell wall would prevent their release to the environment. However, following discovery in fungi, which also have cell walls, EVs have now been described for a variety of Gram-positive bacteria. EVs purified from Gram-positive bacteria are implicated in virulence, toxin release and transference to host cells, eliciting immune responses, and spread of antibiotic resistance. *Listeria monocytogenes* is a Gram-positive bacterium that is the etiological agent of listeriosis. Here we report that *L. monocytogenes* produces EVs with diameter ranging from 20-200 nm, containing the pore-forming toxin listeriolysin O(LLO) and phosphatidylinositol-specific phospholipase C (PI-PLC). Using simultaneous <u>m</u>etabolite, <u>p</u>rotein, and <u>l</u>ipid <u>e</u>xtraction (MPLEx) multi-omics we characterized protein, lipid and metabolite composition of bacterial cells and secreted EVs and found that EVs carry the majority of listerial virulence proteins. Cell-free EV preparations were toxic to the murine macrophage cell line J774.16, in a LLO-dependent manner, evidencing EV biological activity. The deletion of *plcA* increased EV toxicity, suggesting PI-PLC can restrain LLO activity. Using immunogold electron microscopy we detect LLO localization at several organelles within infected human epithelial cells and with high-resolution fluorescence imaging we show that dynamic lipid structures are released from *L. monocytogenes* that colocalize with LLO during infection. Our findings demonstrate that *L. monocytogenes* utilize EVs for toxin release and implicate these structures in mammalian cytotoxicity.

## INTRODUCTION

The pathogenic Gram-positive bacterium *Listeria monocytogenes* is the etiological agent of listeriosis, a disease with serious consequences for pregnant women, newborns, and immunocompromised persons. Healthy individuals who have ingested large amounts of *L. monocytogenes* can suffer from gastroenteritis when the bacterium passes through the gastrointestinal barrier (1-4). *L. monocytogenes* can cause spontaneous abortions in pregnant women and meningoencephalitis by crossing the placental and blood-brain barriers, respectively (5). To invade cells, cross these barriers, and evade the immune system, *L. monocytogenes* has a sophisticated intracellular lifecycle and pathogenic strategy (6, 7).

Initially, *L. monocytogenes* invades various cell types, including non-phagocytic cells, by utilizing two internalins, internalin A (InlA) and internalinB (InlB), with a minor contribution by the pore-forming toxin listeriolysin O (LLO), to induce uptake of the bacterium (1, 5, 8-11). Once internalized in the host vacuole, *L. monocytogenes* employs LLO, phosphatidylcholine-specific phospholipase (PC-PLC) and phosphatidylinositol-specific phospholipase C (PI-PLC) to disrupt the single vacuolar membrane, releasing the bacterium into the cytoplasm (12-16). In the cytoplasm, *L. monocytogenes* replicates rapidly and produces the surface actin assembly-inducing (ActA) protein (17-19). ActA induces actin formation creating a comet tail, ultimately pushing the bacterium towards the host cell surface to invade neighboring cells. In this manner, *L. monocytogenes* replicates and spread within the host avoiding the extracellular space and evading the immune system.

The use of extracellular vesicles (EVs) to secrete compounds to the extracellular space is established in mammals and described in a variety of microorganisms, suggesting these structures are produced by all domains of life (1, 3, 4, 6, 12, 20-24). EVs are small, lipid bilayered spheres ranging in diameter from approximately 20-500 nm. In Gram-negative bacteria, the outer membrane pinches off resulting in the formation of outer membrane vesicles (OMV). OMVs have been associated with, but not limited to, adhesion, immunosuppression, cytotoxicity, virulence, and stress response (20-22, 25, 26) and been postulated to be “virulence bags”. The study of EVs in cell-walled organisms such as Gram-positive bacteria, mycobacteria, and fungi was historically neglected due to the erroneous inference that the combination of a thick cell wall and lack of outer membrane would preclude release of such structures. However, the discovery that fungi produced EVs despite having cell walls (24, 25) stimulated the search for EV in cell-walled organisms. EVs were found in *Bacillus anthracis* and consistent with the idea of “virulence bags”, were implicated in the delivery of anthrax toxin to host cells (12, 17). EV packaging of toxins is widespread: pneumolysin was shown to be released in EVs from *Streptococcus pneumoniae* (4, 27, 28). In *Staphylococcus aureus* and *Mycobacterium ulcerans*, intact toxin-associated EVs are more cytotoxic than disrupted EVs or purified toxin alone, indicating that EV structure is required for efficient delivery of the virulence “package” (3, 29, 30).

There have been two recent reports of EVs in *L. monocytogenes* (17, 31). In this study, we confirmed these findings, extended the observation that EVs are cytotoxic to murine macrophage-like cells, characterized the secreted EVs using simultaneous <u>m</u>etabolite, <u>p</u>rotein, and <u>l</u>ipid <u>e</u>xtraction (MPLEx) multi-omics approach (32, 33), and evidence that EVs are secreted by intracellular bacteria into the cytosol of mammalian cells using electron microscopy and high-resolution fluorescence imaging.

## RESULTS

### L. monocytogenes *produces EVs that carry virulence factors and are hemolytic*

Production of EVs has been demonstrated in several species of Gram-positive bacteria (25). To visualize EVs released from *L. monocytogenes*, we isolated cells or EVs from the extracellular media and imaged these by Transmission Electron Microscopy (TEM). Lipid bilayer vesicles, and vesicle-like structures were visualized protruding from bacterial cells, consistent with EVs (Figure 1A and 1B, see Table 1 for strain description). We tracked the presence of LLO and PI-PLC during our EV purification steps to obtain biochemical evidence for the association of virulence factors, such as LLO and PI-PLC, with EVs (Figure 1C and Supplemental Figure 1). Our data shows that part of LLO and PI-PLC are precipitated by ultracentrifugation at 100,000 *xg*, consistent with partial secretion associated with EVs. Additionally, we performed a centrifugation gradient to discard the possibility that LLO protein aggregates may be precipitating in our assay conditions or that LLO could be co-precipitating with EVs. These assays show LLO density is consistent with EVs and that LLO is protected from protease digestion (Supplemental Fig.1). We explored the temporal relationship between LLO-EV association and bacterial growth by purifying EVs from the wild-type strain at various time intervals during broth culture growth and observed LLO in the EV fraction at all phases of bacterial culture growth (Figure 1D). To ascertain presence of functional LLO in EVs we tested all stages of the purification process for lysis of erythrocytes (Red Blood Cells, RBC) (Figure 1E). The fractions Sup, 100 kDa conc, Sup2, and EVs all lysed RBC. Therefore, we conclude that LLO and PI-PLC are secreted partially in EVs, in agreement with prior reports of EV-associated LLO (17, 31).

**Table 1.** Strains used in this study.

| Strain | Genotype | Phenotype | Source or Reference |
| --- | --- | --- | --- |
| 10403s | wild-type | wild-type | (67) |
| DP-L2161 | 10403s, $\Delta hly$ | LLO<sup>-</sup> | (68) |
| DP-L1936 | 10403s, $\Delta plcAB$ | PI-PLC<sup>-</sup>, PC-PLC<sup>-</sup> | (15) |
| DP-L1552 | 10403s, $\Delta plcA$ | PI-PLC<sup>-</sup> | (34) |
| DP-L1935 | 10403s, $\Delta plcB$ | PC-PLC<sup>-</sup> | (15) |
| GFP | 10403s, GFP | wild-type | (69) |

**Figure 1.**
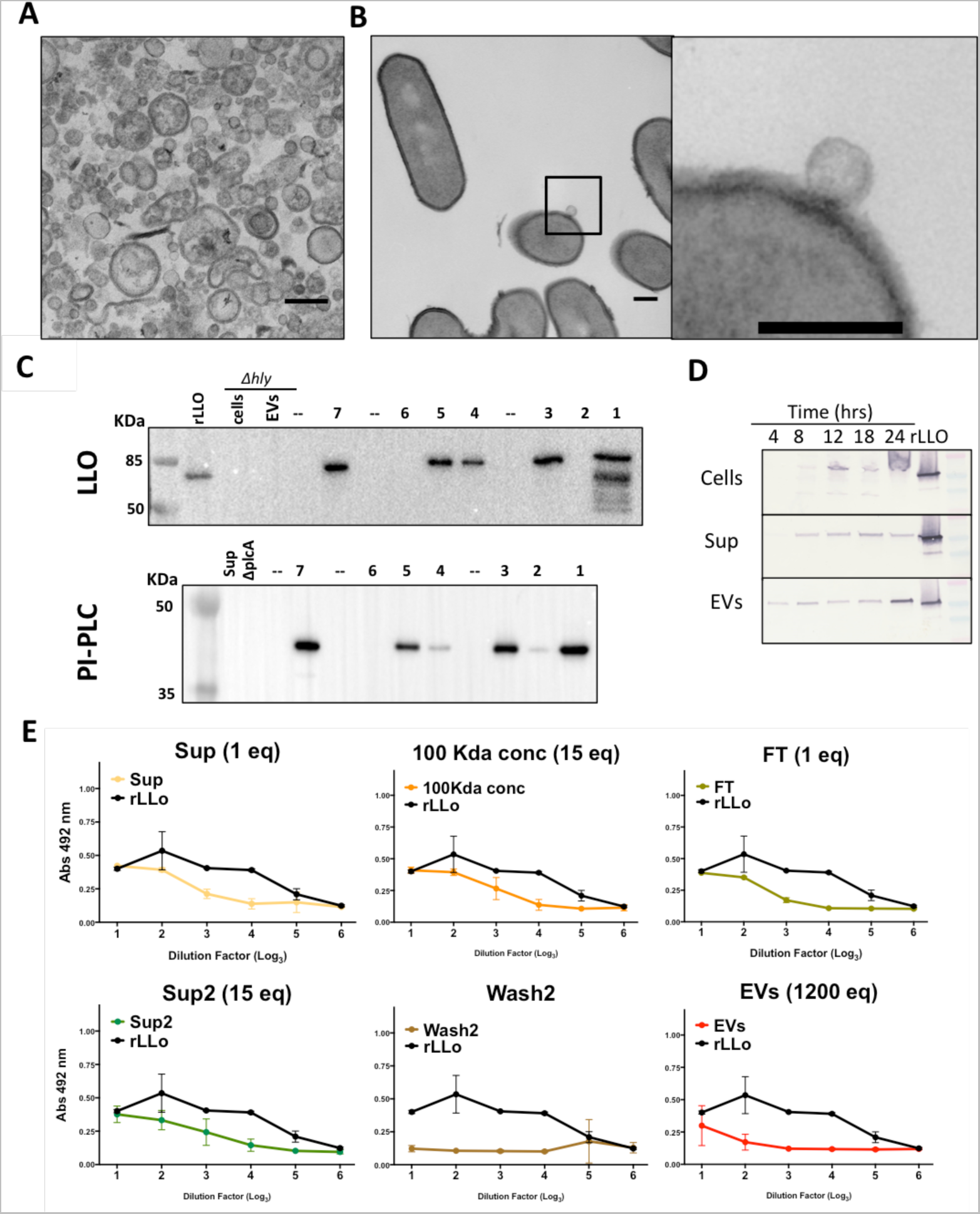
Analysis of EVs from *L. monocytogenes*. EVs were isolated from 18 h culture. (**A**) TEM micrograph of EVs purified from *Δhly* strain culture supernatants. (**B**) TEM micrograph of an EV in the process of release from a *L. monocytogenes* cell. Black arrows indicate putative EV protrusion. Scale bars 200 nm. (**C**) Western Blot of LLO (top) and PI-PLC (bottom) to detect presence during each purification step. Culture supernatants from Δ*hly* (LLO^-^) and Δ*plcA* (PI-PLC^-^) were included as controls. Aliquots were taken at each step and protein extracted by TCA precipitation. Numbers represent fractions from (**Sup. Figure 1** or Materials and Methods); – lanes left empty. Experiments performed twice. (**D**) LLO can be detected in cells, supernatants and vesicles of *L. monocytogenes* from 4 to 24 hr cultures. (**E**) EVs are toxic to RBC. Aliquots of each purification step were serially diluted, added to RBC and released hemoglobin was quantified. Concentration factor relative to cell-free supernatant (Sup) is indicated (1ml of Sup= 1 eq). Experiments were performed twice with two biological replicates in duplicate wells, one representative experiment is shown. FDR 1%. *P<0.05, **P<0.01, ***P<0.001, ****P<0.0001 with oneway Anova. Shown is mean ± SD.

To determine LLO-derived toxicity of EVs in nucleated mammalian cells, J774.16 cells were incubated with EVs purified from cultures of wild-type and avirulent strains Δ*hly* (LLO^-^) and Δ*plcAB* (PI-PLC^-^, PC-PLC^-^) strains, and viability of macrophages was determined based on the MTT assay. EVs purified from the Δ*hly* mutant strain exhibited only residual cytotoxicity, while EVs purified from the Δ*plcAB* strain were more cytotoxic than EVs from wild-type bacteria (Figure 2A). To determine which gene deletion was responsible for increased hemolysis we tested EVs purified from strains with single gene Δ*plcA* and ΔplcB strains. EVs purified from the PI-PLC mutant (Δ*plcA*) were more cytotoxic to macrophages than EVs from other strains. To determine if increased cytotoxicity of EV from Δ*plcA* could be reversed by addition of PI-PLC we performed a rescue assay by combining EVs from the wild-type strain and Δ*plcA* mutant strain, while keeping constant total volume of EVs, and conceivably LLO, added to each sample (Figure 2B). We found that wild-type EVs could partially reverse increased cytotoxicity of Δ*plcA* strain. A similar experiment performed with addition of Δ*hly* EVs to Δ*plcA* EVs confirmed this reversal in cytotoxicity. We investigated whether LLO production was increased in Δ*plcAB* EVs when compared to the wild-type cells but could not detect an increase in LLO secretion (Figure 2C), as shown before for Δ*plcA* (34). To rule out that *ΔplcAB* deletion affected gross composition of EVs we compared EVs from wild-type, *Δhly* and *ΔplcAB* by SDS-PAGE gel analysis. All three strains had similar protein electrophoretic patterns (Figure 2D). The shape and size of EVs from wild-type, *Δhly*, and *ΔplcAB* strains, as measured from TEM micrographs and Dynamic Light Scattering (DLS), were comparable and ranged from 20-200 nm (Figure 2E). The diameter and morphology of EVs from *L. monocytogenes* is similar to EVs produced by *S. aureus* and *S. pneumonia* (1, 3, 4). Hence, the stronger hemolytic action observed for the strain lacking *plcA* was not due to increased production of LLO or EV structural differences.

**Figure 2.**
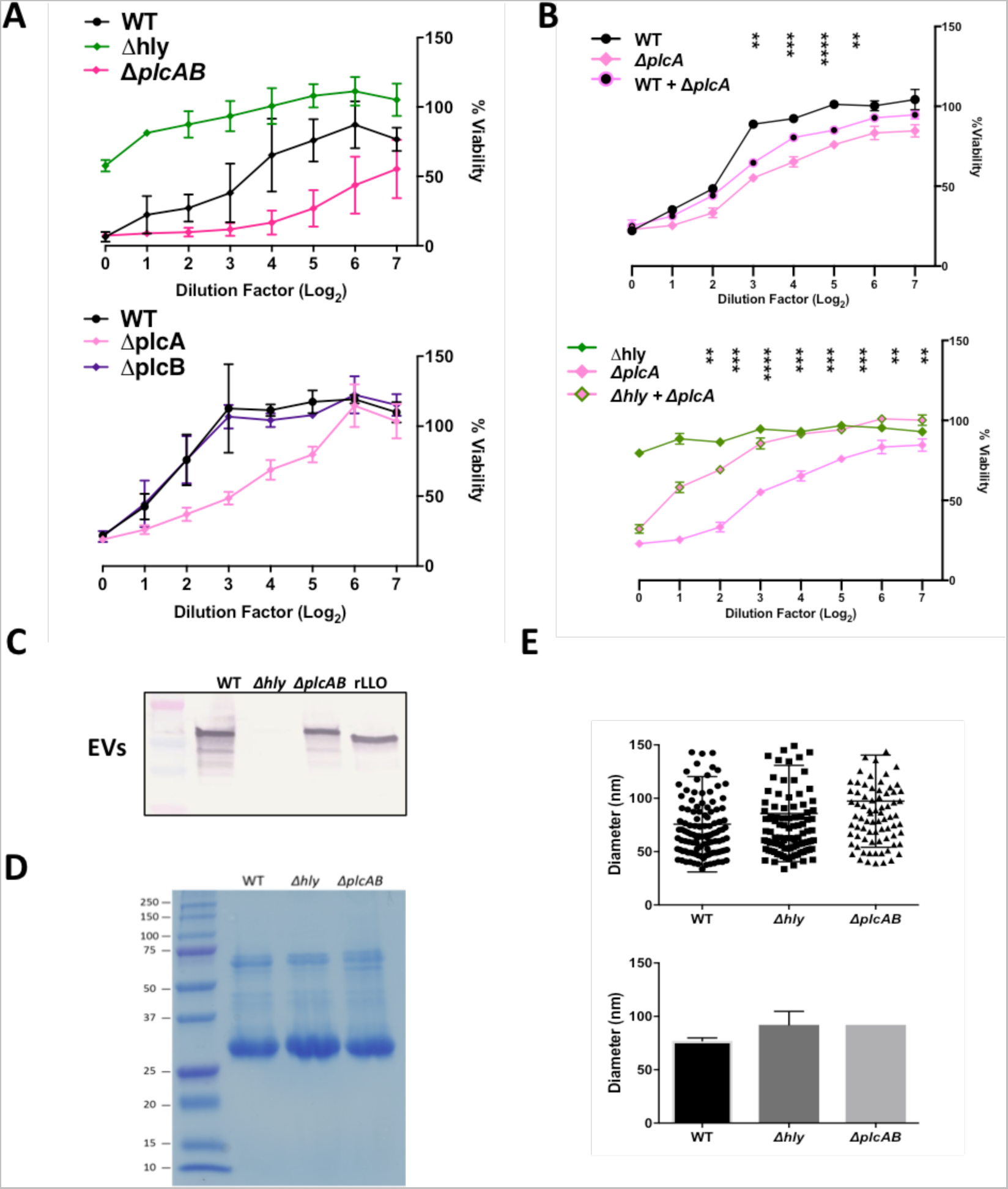
Association between LLO and PI-PLC in EVs. EVs from Δ*plcA* strain were more cytotoxic than wild-type EVs. (**A**) EVs were incubated with J774.16 whose viability was determined by MTT assay. (**B**) Half the volume of EVs from each individual strain was added to maintain equal total volume of EVs. EVs from wild-type and Δ*hly* (LLO^-^) prevent toxicity of Δ*plcA* (PI-PLC) EVs. *P<0.05, **P<0.01, ***P<0.001, ****P<0.0001 with one-way Anova of comparison of Δ*plcA* with the Δ*plcA* + wild-type or Δ*hly*. Experiments were performed once (A and B) in triplicate wells. Shown is mean ± SD. (**C**) LLO content of EVs extracted from wild-type or Δ*plcAB* deletion strains. (**D**) Protein content of EV from *L. monocytogenes* wild-type, *Δhly*, and *ΔplcAB* strains as revealed by Coomasie stained SDS-PAGE. (**E**) EV diameters of wild-type, *Δhly*, and *ΔplcAB* strains of *L. monocytogenes* measured from EVs size based on TEM micrographs and Dynamic light scattering. Shown is average and SD. DLS experiments were performed twice.

### *MPLEx characterization of* L. monocytogenes *cells and EVs*

EVs typically carry proteins, lipids, RNA and metabolites. EVs from *L. monocytogenes* have been characterized for their protein content (17), but no information is available about other components. We performed MPLEx analysis of bacterial cells and EVs to characterize their composition (Fig. 3, Supplemental tables 1 and 2). We found striking differences in protein, lipid and metabolite compositions between cells and EVs. The EVs were enriched in proteins from peptidoglycan synthesis and carbohydrates synthesis (Figures 3A and 3D), suggesting a role in the synthesis of the cell wall. EVs are depleted in proteins associated with translation machinery, and enzymes from fatty acid or amino acid metabolism, but enriched in ABC transporters. We also identified the classical virulence factors of *L.monocytogenes* such as LLO, InlA, InlB, plcB, and ActA associated with EVs, most of them significantly enriched in EVs compared to cells (Figure 3D). Many of the proteins involved in secretion of virulence factors were also detected in EVs (SecDF, SecYEG, SecA, SipZ, YidC, and PrsA2) (35). The lipid profile of bacterial cells showed enrichment in saturated fatty acids and an abundance of phosphatidylglycerol and cardiolipin in bacterial cells, consistent with previous literature (27, 36). Lipid species containing unsaturated fatty acids are more abundant in EVs compared to cells (Figure 3E shows the profile of diacylglycerols as an example). EVs are enriched in phosphatidylethanolamine, sphingolipids (ceramide, monohexosylceramide, mannosylinositolphosphoceramide and sphingomyelin) and triacylglycerols, but depleted in glycoglycerolipids, such as mono and digalactosyldiacylglycerols (Figure 3B and 3E). EVs carried metabolites such as ornithine (or arginine because arginine is converted to ornithine during the sample preparation for GC-MS analysis), citrulline, myo-inositol, phenylalanine, citric acid, and a key intermediate metabolite pyruvic acid (Figure 3C and F).

**Figure 3.**
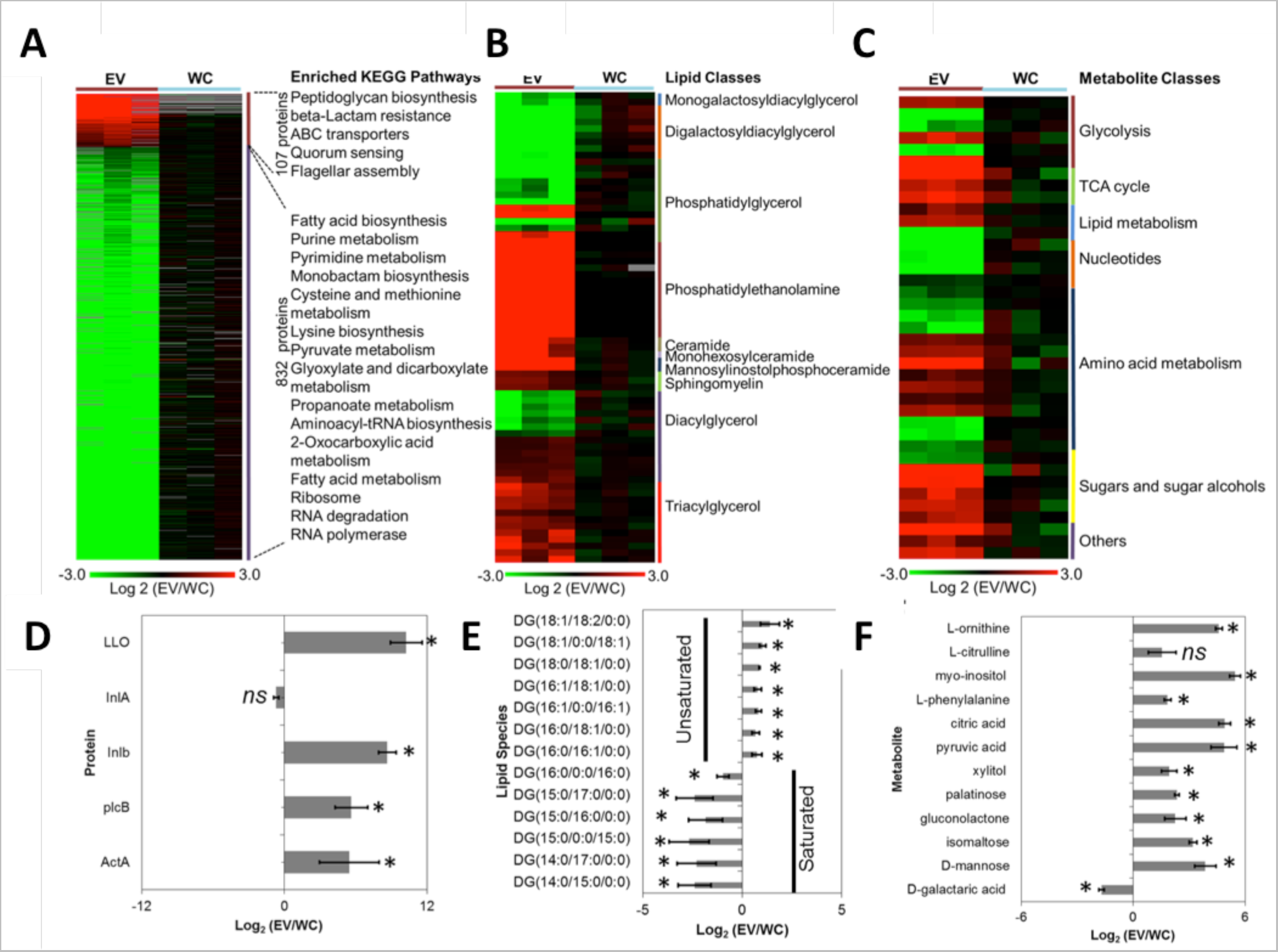
Multiomic analysis of *L. monocytogenes* cells and vesicles. Heatmaps of significantly different (**A**) proteins, (**B**) lipids, and (**C**) and metabolites comparing EV and whole cell (WC) of *L. monocytogenes*. Relative abundances of selected examples of (**D**) proteins, (**E**) lipids, and (**F**) metabolites (**G**). Data represents three independent experiments. *ns* – not significant, * *p*-value ≤ 0.05 for Fisher’s exact test.

### EVs and LLO are produced and target within infected human epithelial cells

We performed immunogold EM (immunoEM) in *Listeria*-infected MCF-7 cells for information on LLO distribution at the ultrastructural level, and in an attempt to visualize association of EVs and LLO in the context of mammalian infection. The specificity of the immunoEM staining using anti-LLO antibodies was determined by assessing LLO-gold particle density. ImmunoEM labeling was negligible for *Δhly* strain compared to wild-type-strain infected cells (Supplemental Figure 2). At 3 hours post-infection (hpi) in MCF7 cells, the vast majority of *Listeria* were detected as free organisms in the host cytoplasm, with only 5% enclosed in a phagosomal compartment (Figure 4A). Some LLO-gold particles were observed in close proximity to free *Listeria* and associated with vesicles, although the origin of these vesicles, if bacterial or the host cell-derived, could not be determined (Figure 4B). Gold particles were abundantly detected in the nucleoplasm of host nuclei, mainly on heterochromatic area (Figure 4C). However, data from immunofluorescence (see below) shows unspecific staining of the mAb to LLO to mammalian cell nuclei, and therefore we advise cautious interpretation of immunodetection when performing studies of LLO location within host cells. A noticeable feature of *Listeria*-infected cells is LLO-mediated induction of autophagy (37). We confirmed the accumulation of autophagosomes and autolysosomes in infected cells, and detected vesicles containing LLO adjacent to autophagic structures (Figure 4D). Additionally, LLO was observed associated with vesicular structures and free aggregates in the host cytosol (Figure 4E-F), without any discernable membrane surrounding or binding the toxin. Another set of host organelles targeted by LLO included tubules of the ER (Figure 4G) and the mitochondria, in which LLO was detected on the outer and inner membranes (Figure 4H). The transport of LLO to host mitochondria may be vesicular as several LLO-containing vesicles were observed surrounding these organelles. The ER or mitochondrial staining was not observed when cells were infected with *Δhly* bacteria. In summary, quantitative distribution of gold particles in mammalian cells containing replicating bacteria reveals that LLO is present at the surface of various host organelles, mainly ER-and mitochondria-associated, and within membrane vesicles whose origin could not determined by immunoEM.

**Figure 4.**
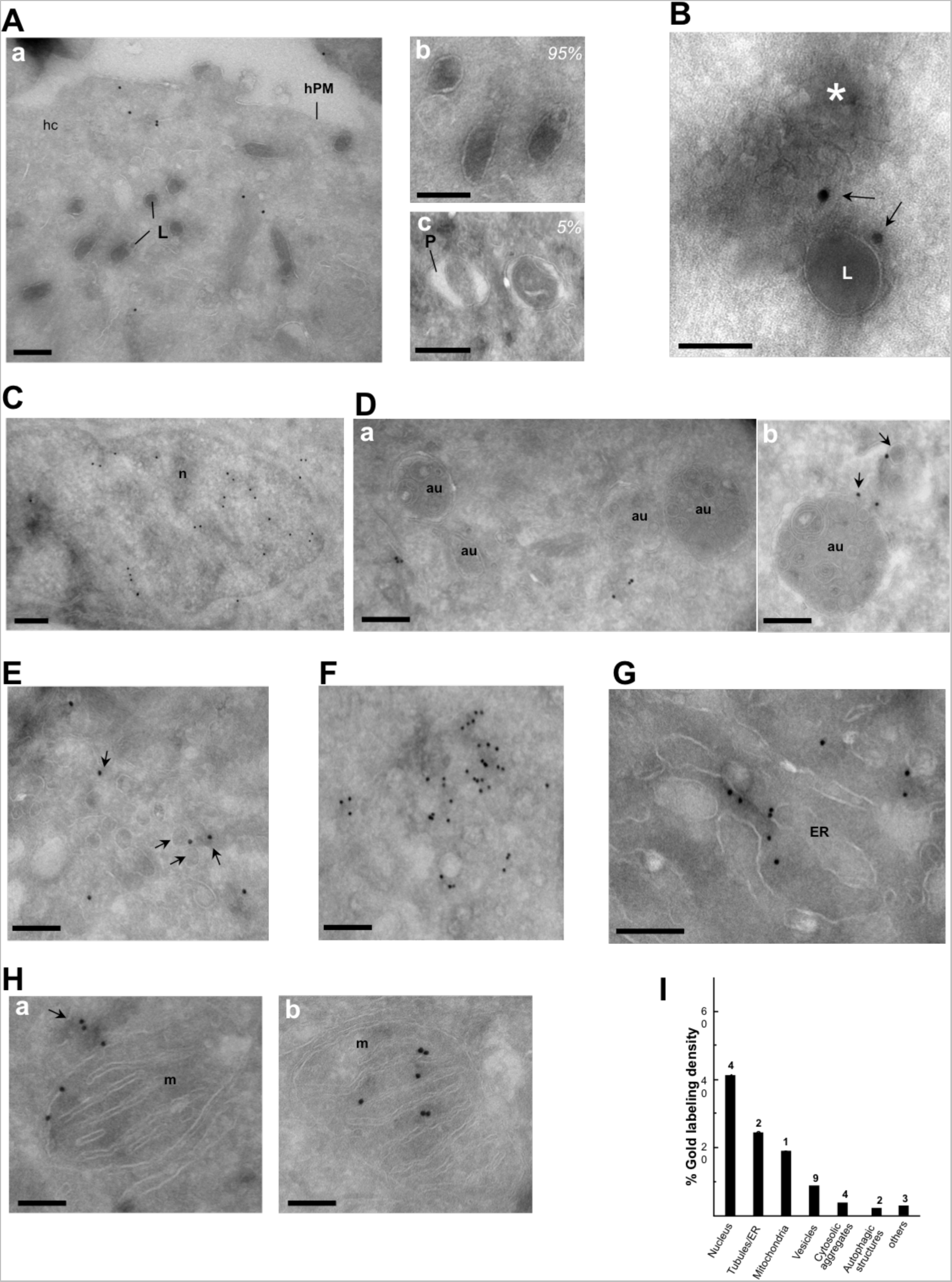
Ultrastructural localization of LLO in *L. monocytogenes*-infected cells. Wild-type (WT) and *Δhly* (LLO^-^) strains of *L. monocytogenes* were grown and used to infect MCF7 cells at a MOI of 20 and at 3 hours post-infection (hpi) cells were fixed and processed for LLO-ImmunoEM. (**A**) Panel a: Large view of infected cell showing free *L. monocytogenes* (L) in the host cell cytoplasm (hc). hPM: host plasma membrane. Panels b and c illustrate free bacteria and intracellular bacteria within phagosomes (P), respectively. (**B**) A cluster of vesicles (asterisk) containing LLO (arrows) in close proximity to *Listeria* (L). (**C**) Host nucleus (n) containing LLO-gold particles. (**D**) Detection of multiple host autophagic profiles (au) in infected cells (panel a), with LLO-stained vesicles in their vicinity (panel b). (**E**) LLO on the limiting membranes of multiple host dispersed cytosolic vesicles (arrows). **F.** Free aggregates of LLO in the host cytosol. (**G-H**) Host ER tubules (G-ER) and mitochondria (m) positively labeled for LLO distributed both on the outer (panel a) and inner membranes (panel b). Note the presence of LLO-positive vesicles close to mitochondria (arrow). All scale bars are 1 μm. (**I**) Stereological analysis of gold labeling demonstrating the specific localizations of LLO in cells infected with *L. monocytogenes*. Density (gold particles per μm^2^) of labeled structures was determined from 55 to 62 cryosections. Percentage of individual intracellular compartment density was determined from the sum of gold density normalized for the variation in distribution of LLO.

As an approach to detect bacterial EV production, dynamics and localization with LLO during mammalian infection we developed a high-resolution imaging approach to detect *Listeria*-derived lipids. To label *L. monocytogenes* lipids, we grew bacteria in the presence of 1 μM BODIPY 558/568 C12 (Bodipy C12), a fluorescent saturated fatty acid analogue (38) for 18 hours. Bacteria cells were grown at 30°C to inhibit the *in vitro* production of virulence genes, including LLO (39), before infecting MCF-7 breast cancer epithelial cells. We note that this may delay progression of infection when comparing to samples imaged by immunoEM. Bodipy C12 effectively labeled bacterial cells (Figure 5 and Supplemental Movie 1), presumably inserted into phosphatidylglycerol-dodecanoic acid lipid species (see lipid analysis in Supplemental Table 2). High-resolution, time-lapse imaging shows that Bodipy C12 puncta were observed both proximal and distal to bacterial cells indicating release and mobility of bacterial lipids, presumably EVs, in the host cytoplasmatic compartment. In another set of experiments, we investigated colocalization of LLO with Bodipy C12-labeled bacterial lipids (Figure 6). At 1.5 hpi, bacteria were mostly localized in phagosomes, Bodipy C12 localized exclusively at GFP-labeled wild-type and LLO^-^ bacteria, while LLO was not detected (data not shown). At 3.5 hpi, LLO was readily detectable as clusters surrounding wild-type bacteria, but not LLO-bacteria (Figure 6A-D). Notably, Bodipy C12 co-localized with LLO puncta at sites proximal and distal, which were not in contact with GFP-labelled wild-type *Listeria* (Figure 6B closed arrows). This data supports LLO association with bacterial lipids and thus compartmentalization within *Listeria*-derived vesicles when infecting mammalian cells. A summary of our results in schematic fashion is shown in Figure 7.

**Figure 5.**
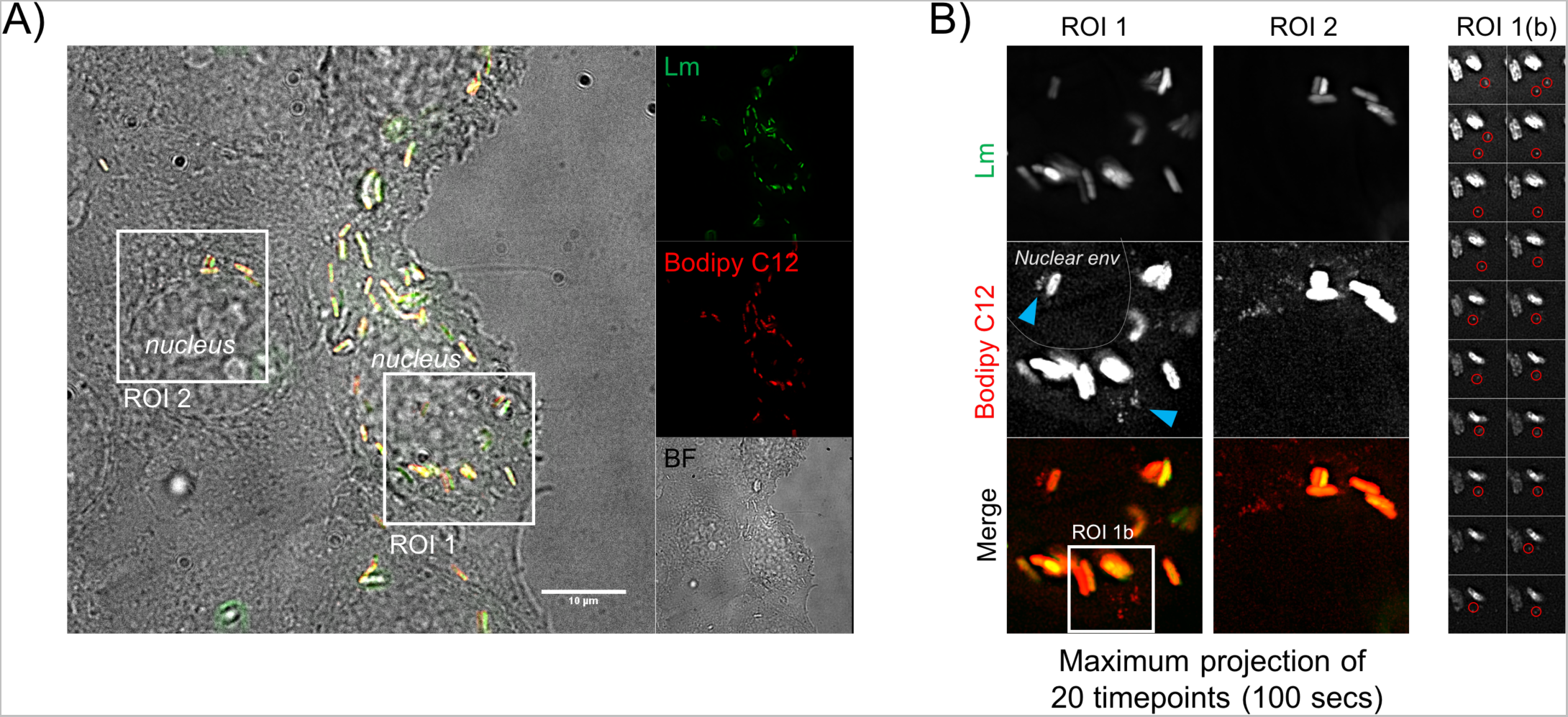
Specific label of bacterial lipids using the fluorescent free fatty acid analog Bodipy C12. GFP-expressing wild-type *L. monocytogenes* were grown overnight at 30°C in the presence of Bodipy C12 (1 μM). MCF7 cells were infected at a MOI of 50 for 30 minutes and 2-3 hpi time-lapse imaging of infected MCF7 cells was performed at 1 image/ 5 seconds. (**A**) Typical infection phenotypes are shown. (**B**) Maximum projection of 20 consecutive images within regions of interest (ROIs) in (A). Clusters of Bodipy C12-positive puncta in the vicinity to *L. monocytogenes* represent the mobility of lipids released from bacteria. (**C**) Mobility of a subset of released bacterial Bodipy C12-positive lipids from ROI 1 (red circles).

**Figure 6.**
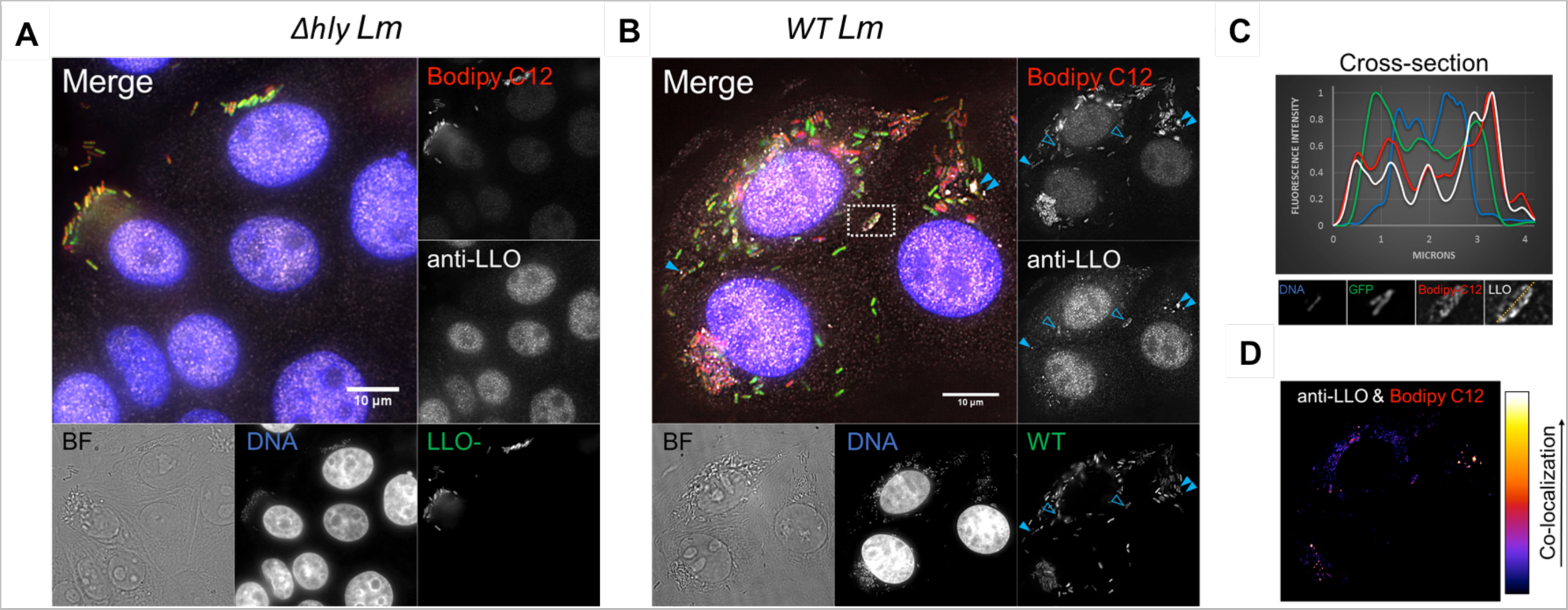
Bodipy C12-labeled lipids *L.monocytogenes* release to cytoplasm and colocalize with LLO. GFP-expressing wild-type (WT) and *Δhly* (LLO^-^) strains of *L. monocytogenes* were grown overnight at 30C in the presence of Bodipy C12 (1 μM). MCF7 cells were infected at a MOI of 50 for 30 min, and at 3 hpi cells were fixed. Fixed cells were immuno-stained using anti-LLO, and DNA was labeled with Hoechst. (**A**) Typical infection phenotypes for *Δhly L. monocytogenes* are shown. Note the anti-LLO labeling was diffuse in the cytoplasm and concentrated in the nucleus. Nuclear staining represent the nonspecific labeling of the LLO antibody, as it was identical for infected and non-infected cells. (**B**) Typical infection phenotypes for wild-type *L. monocytogenes*. LLO is detected in intracellular bacteria (open arrows), and within the cytoplasm (solid arrows). (**C**) Fluorescence profiles for a cross-section of a *L. monocytogenes* with high levels of LLO. (**D**) Co-localization profile for bacterial and cytoplasmic Bodipy C12 and LLO.

**Figure 7.**
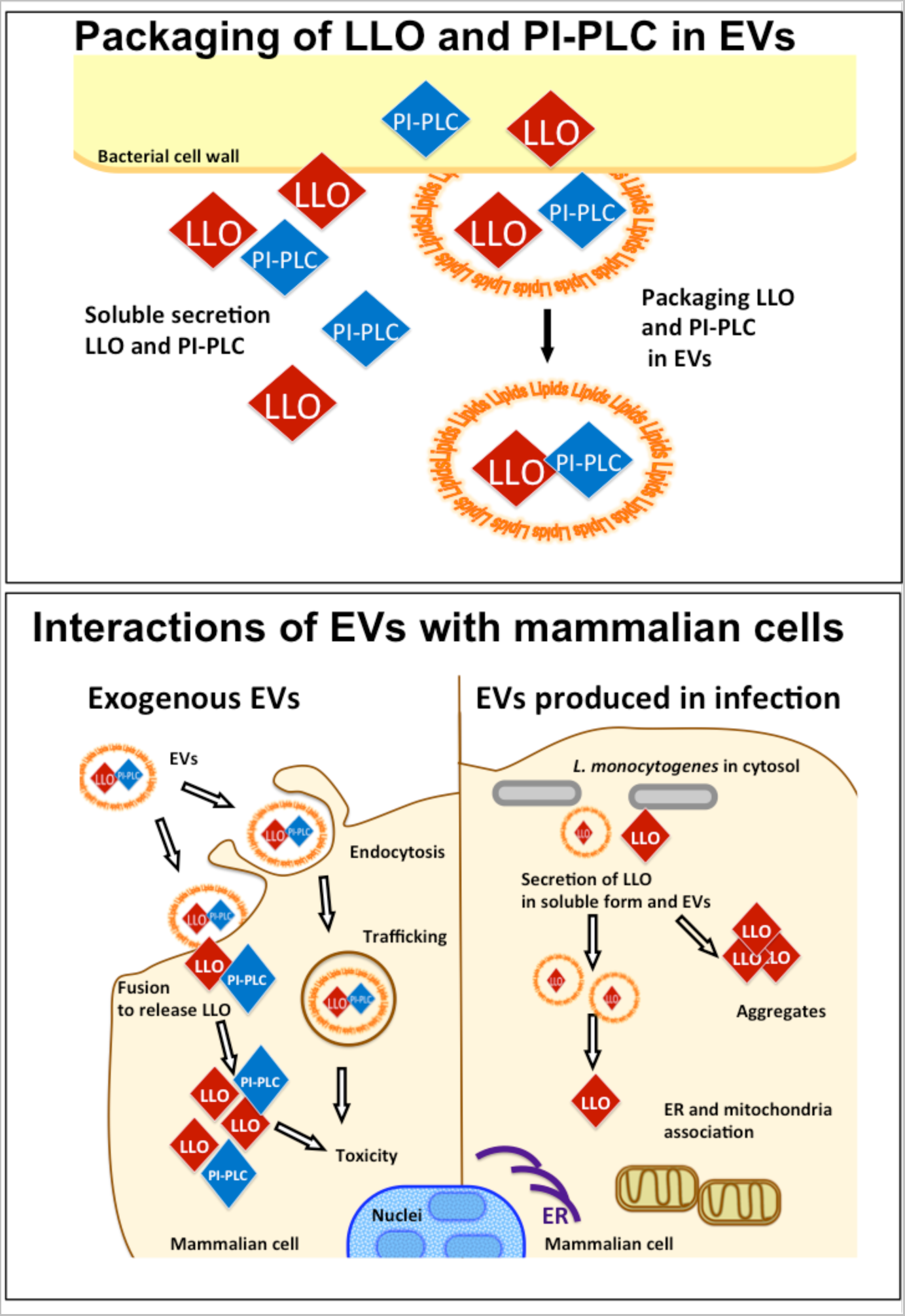
Schematic of major findings of this work. *L. monocytogenes* synthesizes EVs and packages LLO and PI-PLC toxins into EVs. Exogenous EVs interact with, and have significant toxicity towards mammalian cells. Bacterial cells produce EVs as part of their intracellular life cycle and these EVs carry LLO. Secreted LLO into the host cell can remain in EVs or be released in the cytosol, forming large aggregates. LLO interacts with several host organelles (open arrows), likely via its cholesterol-binding properties.

## DISCUSSION

The production of EVs has now been described in all domains of life. OMVs from Gram-negative bacteria have been studied for over half a century, while only recently EVs were found in Gram-positive bacteria as diverse as *Staphylococcus aureus* (1), *Bacillus anthracis* (12), *Bifidobacterium spp*. and *Lactobacillus spp*. (40), *Bacillus subtilis* (23), *Pneumococcus pneumoniae* (4), *Streptomyces lividans* (41), *Clostridium perfringens* (20) and *Listeria monocytogenes* (17, 31). One advantage for toxin packaging in EVs is that it allows their delivery as a concentrated warhead that is not diluted as a function of the square of the distance from the bacterial surface. The field of bacterial EVs has historically been plagued by controversy as to whether these structures were real cellular products or the result of lipid association following lysis of bacteria. For Gram-positive bacteria these questions are further complicated by concerns as to how EV traverse the bacterial cell wall (23). Raising additional suspicion for their existence was the absence of EV-null mutants, although recent work have identified genes modulating EV synthesis in Gram-positive bacteria (42) and mycobacteria (43), and gene deletions that alter morphology of EVs (17). In this study we establish that EVs are associated with LLO via biochemical purification approaches, functional detection, immunoEM and fluorescence microscopy. We provide evidence against their formation as by-products of cell autolysis by showing major differences in lipid and protein composition from parent bacteria and imaging their release inside infected mammalian cells. These findings together with the two reports (17, 31) provide compelling evidence for the notion that *L. monocytogenes* releases EV packed with virulence factors.

When establishing toxicity of EVs for nucleated mammalian cells we detected increased cytotoxicity in EVs from strains deleted in *ΔplcA* (but not ΔplcB deletion). These results are surprising since no alteration of hemolysis capacity was observed upon engineering of the Δ*plcA* DP-L1552 strain (34). One explanation is to attribute these discrepancies to different experimental models: source of toxins (EVs versus supernatants) and assays (RBC hemolysis vs cytotoxicity assays). However, we entertain other possibilities that could account for these findings. Firstly, it is possible that *ΔplcA* loss subtly affects EVs into more efficient vehicles of LLO delivery since we could not detect gross structural abnormalities relative to wild-type strains. Secondly, it is possible that PI-PLC interacts with LLO to reduce its toxicity in some manner (43), but the mechanism by which this phenomenon occurs is unclear at this time.

Overall, our data shows that EVs have distinct compositions from the bacterial cells, strong evidence that EVs are specialized structures from the bacterium. Exosomes from mammalian cells have distinct a lipid composition cellular membrane of parent cell, with exossomes enriched in sphingomyelin, and cholesterol (29). The saturated lipids in *L. monocytogenes* cells are necessary in low temperature conditions (27, 28). EVs are enriched in sphingolipids, phosphatidylethanolamine, with a higher component of unsaturated fatty acids. Remarkably their compositions were distinct from other bacterial EVs: EVs from Group A *Streptococcus* (42) displayed an enrichment in phosphatidylcholine and monoacylglycerol. Lipid composition of EVs may vary with the bacterial species and possibly is finely adapted for each environmental niche. We note that our experimental approach is not suitable to detect steroids lipids, such as bacterial hopanoids or mammalian cholesterol, but the importance of steroids in lipid bilayer stability warrants a detailed study of the steroid lipid class in EVs. The same pattern emerged for detection of metabolites: EVs are strikingly different from bacterial cells. Both lipids and metabolites are still quite mysterious components of EVs and further inferences are precluded by the fact that characterization of EVs lipids and metabolites is still quite rare.

EVs carry significant variety of toxins for the parent bacterium, in particular LLO (17, 31). We note prior studies that recovered LLO from culture supernatant through protein purification would have included the vesicular fraction and therefore those results are compatible with our findings (44). Our findings do not rule out that LLO is released at the bacterial surface in its free form, but suggest instead simultaneous soluble and vesicular secretion. Redundancy of secretion mechanisms is consistent with the fact that strains deleted for factors involved in secretion and folding of LLO retain some hemolytic activity: both PrsA2 (45) and SecDF (46) deletion strains possess 30% hemolysis capacity compared to wild-type bacteria. Further SecDF, PrsA2 and SipZ were detected in EVs, suggesting that these components may be involved in both soluble secretion and EV-mediated secretion (47), i.e., the pathways overlap such that the machinery involved in secretion of the soluble form is equally involved in secretion of the vesicular form. Dual secretion is not the rule for EV-associated toxins. For example in *B. anthracis* the majority of anthrax toxin was secreted via EVs, since toxin was present in the EV pellet and not the supernatant after ultracentrifugation (12). It remains to be seen how sorting of the soluble vs vesicular form occurs. The EVs are enriched in proteins from peptidoglycan synthesis as well as carbohydrates, suggesting EVs are involved in the synthesis of the cell wall. It is conceivable that some of these enzymes may be involved in biofilm formation (32). However, we could not detect in our EV protein dataset any of the enzymes reported to be involved in biofilms of *L. monocytogenes*, but this may just be explained due to our use of non-biofilm inducing conditions.

Another intriguing aspect is whether exogenously added EVs exert their toxicity by fusing with host plasma membrane and release of contents into the host or instead EVs are first ingested by the host and trafficking in the endocytic/phagocytic compartments precedes release of toxins (31). In other bacterial pathogens (*S. aureus* and *Legionella pneumophila*), EVs were found to fuse with host cells (48) but it has been suggested that EVs from *L. monocytogenes* are internalized in endosomes (31). In the absence of a detailed study we consider it is possible that EVs from *L. monocytogenes* may simply be uptaken by endocytosis, or, given the invasive arsenal of *L. monocytogenes*, equally probable that EVs induce ingestion by host cells, either via internalin (InlA and InlB) interactions with lipid microdomains at the cell surface (49), or by LLO pore-triggered entry mechanisms analogous to what has been observed for *L. monocytogenes* cells invading human hepatocytes (9).

The majority of previous studies have focused on exogenous addition of EVs and its effects on host cells (31), but unequivocal demonstration of EV secretion within infected host cells is e demosntrated. Two reports have described synthesis of EVs within murine and human macrophages by *M. tuberculosis* (52) and *L. pneumophila* (53) because infected macrophages cells secreted disparate populations of EVs with one population carrying solely host proteins while the second population carried mostly bacterial products. We observed LLO associated with small vesicles whose size was compatible with that of EVs by immunoEM of mammalian cells infected with *L. monocytogenes* (150-250 nm in diameter in infection versus 90nm measured by DLS). An alternate explanation for this finding is that LLO molecules could hijack mammalian lipids via interaction with cholesterol, the only known mammalian cell receptor for LLO (50). Regardless of the origin of these vesicles, either via bacterial EVs or hijack of host membrane, their proximity to host organelles suggests a vesicular delivery of LLO, preceding insertion into organelle membranes. Fluorescence microscopy demonstrated bacterial lipids distal to bacterial cells within host cytosol and bacterial lipid colocalization with LLO supporting lipid-EV association with LLO and therefore suggesting that even in host cytosol LLO-EV association exists. In the future, strategies akin to ours may allow to observe EVs contribution to pathogenesis in real time and one may even identify and tag a lipid species that is inserted solely in EVs facilitating a finer distinction from bacterial cell-associated lipids. LLO, “the swiss-army knife” (8), is continuously produced to regulate multiple steps of the intracellular life of *L. monocytogenes*. LLO secreted by extracellular *L. monocytogenes* or addition of recombinant LLO to mammalian cells affects the ER (55) and mitochondria (51), interaction of LLO with cholesterol causes small membrane perturbations *in situ* and the damaged cellular compartments act as intracellular danger signals, triggering autophagy (57) such that tight control of toxin synthesis and activity is needed to avoid killing the host cell (52). Future studies will determine how LLO compartmentalized within EVs can incorporate this paradigm (53). The functions of intracellular EVs, and their LLO cargo, may simply recapitulate functions of soluble LLO such that their roles are indistinguishable or, alternatively, vesicular LLO could facilitate a particular set of interactions. For example in *B. anthracis* EV integrity was required for toxicity of EV-toxin cargo (12).

In summary, we confirm that *L. monocytogenes* EV contain LLO, as well as many other virulence factors. *L. monocytogenes* provides yet another example of a Gram-positive bacterium packaging its toxins in EVs (17, 24, 40). The fact that many Gram-positive and cell-walled microorganisms produce EVs raises fascinating questions in cell biology regarding the mechanisms for transit across cell walls and the packaging of EV cargo. Our results confirm *L. monocytogenes* produces EVs, characterize composition of EVs and establish LLO-EV association during mammalian infection. Our findings provide fertile ground for future investigations in this rapidly developing field.

## EXPERIMENTAL PROCEDURES

### Bacterial strains and mammalian cell lines

*L. monocytogenes* strains were maintained at −80°C (Table 1). All cultures were grown in Brain Heart Infusion (BHI) broth (Difco) with 180 rpm shaking at 37°C, except when otherwise noted. J774.16 macrophage-like cells were maintained in 10% fetal bovine serum, 10% NCTC, 1% non-essential amino acids, in Dulbecco’s Modification of Eagle’s Medium (DMEM catalog #10-013-CV). Human MCF-7 breast cancer cells (ATCC HTB22) were maintained in DMEM, supplemented with 10% FBS, L-glutamine, non-essential amino acids, and penicillin/streptomycin/amphotericin B (all from Thermo Scientific).

### EVs purification

Cultures were incubated at 180 rpm shaking at 37°C for 18 h of growth unless specified otherwise. EVs were purified from planktonic BHI cultures (see diagram on Supplemental Figure 1). Cultures were centrifuged at 16,000*xg* for 15-20 min to remove cells and subsequently filtered through a 0.2μm filter to remove remaining cells and debris. Cell-free supernatant (“Supernatant”-#1) was concentrated using a 100 kDa cutoff membrane on an Amicon ultrafiltration device (Millipore). In some experiments the fraction that was able to flow through the membrane was collected (“Flow-through”-#2). The fraction above the membrane ((“100 KDa Conc”-#3) was washed with PBS (with Ca^2+^ and Mg^2+^), brought down to 10-50 mL while the flow through collected as a separate fraction (“PBS Wash 1”-#4). About 10-50 mL remaining on the top of the membrane were then ultracentrifuged at 100,000 xg for 1h in a Beckman Optima-XL to pellet EVs. Supernatant was removed (“Supernatant 2”-#5), washed once with PBS and re-centrifuged. At the end of this centrifugation, supernatant (“PBS Wash 2”-#6) was collected and pellet (“EVs”-#7) was resuspended in PBS. EVs were to gradient centrifugation ranging from 45 to 10% Optiprep (Sigma-Aldrich) in HEPES buffer, as previously described (54, 55). For protection assays the 100KDa conc was subject to 0.1% SDS, pH adjusted to 7.4 or treated with trypsin 0.005% (Corning) at 180 rpm shaking for 1 h at 37°C, followed by regular purification of EVs. For this experiment Sup2 was concentrated using 10 KDa Centricon. Protease inhibitors (cOmplete, Roche) were added to all samples after ultracentrifugation steps or after completion of TCA extraction.

### Transmission Electron Microscopy

Cell and EV samples for TEM were processed as previously described (23, 44). Samples were viewed on a JEOL 100CXII or JEOL 1200EX at 80kV. For measurements of EV diameter by TEM a 1.27 correction factor was applied to compensate for the fact that EM diameters are derived from spheres (56).

### Immunodetection of LLO

For visualization of proteins from bacterial cells, approximately 100 mL of culture was spun to obtain bacterial and these cells were disrupted by sonication (4 rounds of 20 s sonication and 30 s rest on ice) (Sonic Dismembrator Model 100; Fisher Scientific). Supernatants were isolated from 1 L cultures, concentrated using a 100-kDa ultrafiltration and then further concentrated on a 30-kDa ultrafiltration device (Amicon, Millipore). The equivalent of 1 mL cells, and 1 L of EVs were supplemented with 4x Sample Buffer and 20μl used for each sample. All buffers used with intact cells and EVs were at mammalian physiological pH (7.0-7.4). Culture BHI broth after overnight (18 h) culture had pH=6.3. Samples were kept on ice at all times, except ultrafiltration was performed at room temperature. Samples were then incubated with 15% TCA with 1/100 of its volume of 2% DOC or with 10% TCA overnight at 4°C. The resulting pellet was washed 2 times with acetone, resuspended in protein sample buffer (50mM Tris, 2% SDS, 100mM DTT, 10% glycerol) in a volume proportional to the equivalent starting culture material (l00 μL per 1000 mL of starting culture)., Equal volumes of samples were run on a denaturing 10% Bis-Tris gel (Nupage, Life Technologies), according to manufacturer instructions. Alternatively samples were spotted onto a PVDF membrane using a dot blot apparatus. Positive control was recombinant ~53kDa LLO fragment (Abcam #ab83345) and negative control was supernatant from deletion strain. PI-PLC rabbit antiserum was a kind gift of Dr. Howard Goldfine (57). Samples were transferred into a PVDF membrane and immunoblotted using anti-LLO polyclonal rabbit IgG (Abcam #ab43018, lotGR56599-8, 17, or #ab200538 lotGR211194-7,12) in blocking buffer (5% non-fat dry milk in TBS-1% Tween) overnight at 4°C. Secondary antibodies goat α-rabbit IgG-HRP (Southern Biotech) was used for 1 h at room temperature. Blot was developed using SuperSignal West Pico Chemoluminescent Substrate (Thermo Scientific). For visualizing proteins, gels were incubated at room temperature in Coomassie Brilliant Blue R-250 (Thermo Scientific) for 1 h and then destained with water, methanol, and acetic acid (50/40/10 v/v/v) overnight.

### Erythrocyte (RBC) lysis assay

Erythrocyte (red blood cells, RBC) lysis was measured as described previously. Briefly sheep RBC (Innovative Research) were resuspended in 1% BSA with 5 mM DTT at 5% suspension. Aliquots from each purification step were diluted serially in a 1:3 ratio. Assay was performed in total volume of 300 μl and concentration of rLLO of 350 ng/mL. EVs (obtained as described above) were resuspended in PBS or 10 mM dextrose, filter sterilized, and diluted in 1%BSA. Samples were incubated with RBC for 30 min at 37°C, spun briefly to sediment intact RBC, and absorbance of the supernatant was measured at 594 nm.

### Cytotoxicity MTT Assay in J774.16 cells

MTT [3,(4,5-dimethylthiazol-2-yl)2,5-diphenyltetrazolium bromide] colorimetric assay was utilized to determine EV cytotoxicity to J774 murine macrophages. EVs were purified from 1 or 2 L of broth culture and filter sterilized with 0.22 μm. A 96-well plate was seeded with 5x104macrophages per well and incubated overnight at 37°C. EVs were resuspended in PBS 10% dextrose, filter sterilized and added to cells in triplicate with a serial dilution of 1:2 in DMEM. The plate was incubated for 4 h at 37°C. At the end of 4h, sterile MTT solution (5 mg/ml in PBS) was added to wells and incubated for an additional 2 h at 37°C. A purple color forms due to reduction of MTT to formazan by viable macrophages. Formazan precipitate was dissolved overnight at 37°C upon addition of extraction buffer (12.5% SDS, 45% dimethylformamide in PBS). Absorbance was read at 570 nm. EV killing was compared to negative control of 10% dextrose (same volume as EVs) in media and 100-200 ng of purified LLO fragment (Abcam#ab68200). When treating J774.16 cells with EVs from two strains, the total volume of EVs was maintained by adding 50% the volume of EVs from each individual strain.

### Dynamic Light Scattering

Hydrodynamic diameters of EVs were measured with 90Plus/BI-MAS Multi Angle Particle Sizing analyzer (Brookhaven Instruments Corp.), as described previously (23, 58).

### Multi-omics experimental design and statistical rationale

For the multi-omics experiment *Listeria monocytogenes* cells were grown and harvested in biological triplicates, which is enough for studying samples derived from established cell cultures (63). Furthermore, we observed that with the recent improvements of the mass spectrometers the technical variability is much smaller than natural biological variability, avoiding the necessity of collecting data for technical replicates. To ensure the performance of the mass spectrometer we run quality control samples before and after each sample batch, and quality is monitored as previously described (64). All the samples were randomized for sample preparation and again for data collection. Enrichment or repletion of molecules in EVs was determined by T-test considering two-tailed distribution and equal variance.

### Metabolite, Protein and Lipid Extraction (MPLEx)

*L. monocytogenes* cells were lysed by vigorous shaking with 0.1 mm zirconia/silica beads as in 50 mM NH_4_HCO_3_ buffer, pH 7.8. Then cell lysates and EVs were submitted to MPLEx, as previously described (33). Briefly, 5 volumes of −20°C chloroform:methanol (2:1) were added and the samples incubated for 5 minutes on ice, before vortexing for 1 minute and centrifuging at 12,000 rpm at 4°C for 10 minutes. The top and bottom phases, containing metabolites and lipids respectively, were collected into autosampler vials and dried in a vacuum centrifuge (Labconco, Kansas City, MO). The protein pellet was washed by adding 1 mL −20°C methanol and centrifuging at 12,000 rpm for 10 minutes at 4°C. Then the supernatant was discarded and precipitated protein was dried in a vacuum centrifuge.

### Proteomic analysis

Proteins were digested with trypsin as described elsewhere (33) and resulting peptides were analyzed by liquid chromatography tandem mass spectrometry (LC-MS/MS) in NanoAcquity UPLC (Waters) connected to a Q-Exactive mass spectrometer (Thermo Fisher Scientific). Peptides were loaded into trap column (5 cm x 360 μm OD x 150 μm ID fused silica capillary tubing, Polymicro, Phoenix, AZ; packed with 3.6-μm Aeries C18 particles, Phenomenex, Torrence, CA) and gradient was performed in a capillary column (70 cm x 360 μm OD x 75 μm ID packed with 3-μm Jupiter C18 stationary phase, Phenomenex): 1-8% solvent B in 2 min, 8-12% B in 18 min, 12-30% B for 55 min, 30-45% B in 22 min, 45-95% B in 3 min, hold for 5 min in 95% B and returning to 1% B in 10 min. Solvent A: water with 0.1% formic acid and solvent B: acetonitrile (ACN) containing 0.1% formic acid. Spectra were collected in 400-2000 m/z range with a resolution of 35,000 at m/z 400. MS/MS was performed once on the top 12 most intense ions with ≥ 2 charges using high-collision energy (HCD) (30% NCE and resolution of 17,000 at m/z 400) and dynamic exclusion was set to 30 s. Spectra were queried against the *Listeria monocytogenes* 10304s protein sequences from Uniprot Knowledgebase (2815 sequences downloaded on January 17, 2017) using MaxQuant (v.1.5.5.1) (59), considering only fully tryptic peptides with two missed cleavages allowed, methionine oxidation as a variable modification and cysteine carbamidomethylation as fixed modification. Peptide mass tolerance was set at 20 ppm first database search and at 4.5 ppm after calibration for the main peptide search. Fragment mass tolerance was set at 20 ppm for both searching rounds. The peptide score threshold was set as the default parameter from MaxQuant and the FDR was limited to 1% in both peptide-spectrum match and protein levels. Quantification was performed using the LFQ function of MaxQuant using a minimum ratio count of 2, minimum number of neighbors of 3 and average number of neighbors of 6. Redundant peptides were assembled into protein groups and quantification of proteins was performed based on unique peptides plus razors. Extracted LFQ intensities were submitted to linear regression and central tendency normalization with InfernoRDN (formely DAnTE) (60). Function-enrichment analysis was performed using the KEGG annotation (61) and considering only pathways with more than 3 entries, fold enrichment ≥ 1.5 and p-value ≤ 0.05 (Fisher’s exact test).

### Metabolomic analysis

Polar metabolites from both the EVs and cell lysates were derivatized and analyzed as described previously (33) using a GC 7890A GC-MS system (Agilent Technologies). Blanks and fatty acid methyl ester (FAME) samples were included in the analyses for background reference and RT calibration purposes, respectively. GC-MS data was processed Metabolite Detector as previously described (62). For identification purposes metabolites were identified by matching experimental spectra to a PNNL augmented version of FiehnLib library (63), containing spectra and validated retention indices of more than 900 metabolites. As for the unidentified metabolites, these were screened against the NIST14 GC-MS Spectral Library by comparing their spectra alone (denoted with “NIST”). The curated data set of identified metabolites, unidentified features, and their abundances for each sample was then subjected to multivariate data analysis (MVDA) by making use of MetaboAnalyst (64).

### Lipidomic analysis

Lipids were analyzed by LC-MS/MS both positive and negative ionization modes in a LTQ Obtrap Velos mass spectrometer (Thermo Fisher Scientific) as described in detail previously (65). Lipid species were identified using the LIQUID tool (65) followed by manual inspection. Confidently identified lipid species were quantified using MZmine 2 (66) and the peak intensities were normalized by linear regression and central tendency with InfernoRDN.

### Immunoelectron microscopy of L. monocytogenes infected MCF-7 cells

Cell lines were plated at 1x10^5^/mL the day before infection in 2 ml for 6 well plates. *L. monocytogenes* cells from an overnight starter culture in BHI 37°C were inoculated in fresh media at 1/100 dilution and grown for 1-2 h in the same conditions. Monolayers of MCF-7 cells were infected with *L. monocytogenes*, wild-type or LLO^-^(*Δhly*), at the MOI of 1:20 for 3 h before fixation, first in 4% paraformaldehyde (PFA; Electron Microscopy Sciences, PA) in 0.25 M HEPES (pH=7.4) for 1 h at room temperature, then in 8% PFA in the same buffer overnight at 4oC. Samples were infiltrated, frozen and sectioned as previously described (73). The sections were immunolabeled with mouse anti-rabbit LLO antibody (Abcam #ab200538) at 1/20 dilution in PBS/1% fish skin gelatin, then with IgG antibodies, followed directly by 12 nm protein A-gold particles before examination with a Philips CM120 Electron Microscope (Eindhoven, the Netherlands) under 80 kV.

### Immunofluorescence studies of L. monocytogenes-infected MCF-7 cells

MCF-7 cells were plated and infected in 8-well microscopy μ-slides (ibidi, Germany) in antibiotic-free media. To label *L. monocytogenes* membrane lipids, bacteria were incubated with 1 μM Bodipy 558/568 C12 (Thermo Fisher Scientific) for 18 h at 30°C, to repress LLO expression, and excess dye washed off prior infection of mammalian cells at an MOI of 1:50. Gentamicin was added 30 min after start of infection to repress extracellular growth of bacteria. For time-lapse imaging wild-type GFP-expressing *L. monocytogenes* was used and imaged 2-3 hpi hpi, in a humidified chamber at 37°C with 5% CO_2_. For immunofluorescence staining, cells were fixed for 15 min with 4% paraformaldehyde (in PBS, pH 7.4) at the indicated time points. Cells were then permeabilized with 0.3% Triton X-100 in PBS, blocked with 3% BSA, and incubated with primary antibodies against LLO (Abcam; #ab200538) overnight at 4°C. Staining was performed for 30 min at room temperature using hig*hly* cross-absorbed Alexa Fluor 647 secondary antibodies (Thermo Fisher Scientific). DNA was labeled using Hoechst 33342 (1ug/mL; Thermo Fisher Scientific). 3D stack of infected cells were obtained using a DeltaVision Elite microscope system (GE Healthcare) equipped with a 60x oil immersion objective. Images were deconvolved (SoftWoRx) and analyzed using ImageJ (https://imagej.nih.gov/ij/). Colocalization was performed using the Colormap plugin (74).

### Statistics

Error bars represent mean and SD. Statistics tests were performed using GraphPad Prism. Diameters were analyzed by one-way ANOVA with Bonferroni correction.

## ACKNOWLEDGEMENTS

AC was supported by National Institutes of Health awards 5R01HL059842, 5R01AI033774, 5R37AI033142, and 5R01AI052733. The MPLEx experiment was performed in the Environmental Molecular Science Laboratory, a U.S. Department of Energy (DOE) national scientific user facility at Pacific Northwest National Laboratory (PNNL) in Richland, WA. Battelle operates PNNL for the DOE under contract DE-AC05-76RLO01830. Multi-omics data were deposited in the MassIVE repository under accession numbers MSV000081402, MSV000081403 and MSV000081404.

We thank Geoffrey Perumal, Benjamin Clark, and Leslie Gunther at Analytical Imaging Facility, Albert Einstein College of Medicine for assistance with the transmission electron microscopy. We would like also to thank the technical competence of Kimberley Zichichi from the Electron Microscopy Facility at Yale University for the immunoEM. We thank Howard Goldfine, University of Pennsylvania, for the kind gift of PI-PLC antiserum. The data in this paper were from a thesis that was submitted by L.B. in partial fulfillment of the requirements for the degree of Doctor of Philosophy in the Sue Golding Graduate Division of Medical Science, Albert Einstein College of Medicine, Yeshiva University, Bronx NY.

## Author Contributions

LB, CC, MM and RPR performed EV isolation, and experiments. AHB and NRB performed IF, IC performed IEM. ESN designed the MPLEx experiments and analyzed the data, MCB performed proteomic analysis, JEK the lipidomic analysis, and HMH the metabolomic analysis. GL supplied bacterial strains and critical scientific advice. LB, CC, IC, NRB and AHB and AC designed the experiments and wrote the manuscript. All authors read and approved the final version of the manuscript.

## Supplemental Files

**Coelho_etal_MPLEx_data_SupplementalTables.xlsx**

Supplemental Table 1. Proteomics-MPLEx data *L. monocytogenes* EVs (identified peptides).xlsx

Supplemental Table 2. Lipidomics-MPLEx data *L. monocytogenes* EVs (protein groups and quantification).xls

Supplemental Table 3. Lipidomics-MPLEx data *L. monocytogenes* EVs.xls

Supplemental Table 4. Metabolomics-MPLEx data *L. monocytogenes* EVs.xls

**Supplemental_movie 1. Time-lapse imaging of *L. monocytogenes* lipid’s mobility within cytosol of MCF-7 cells.**

**Supplemental Figure 1.**
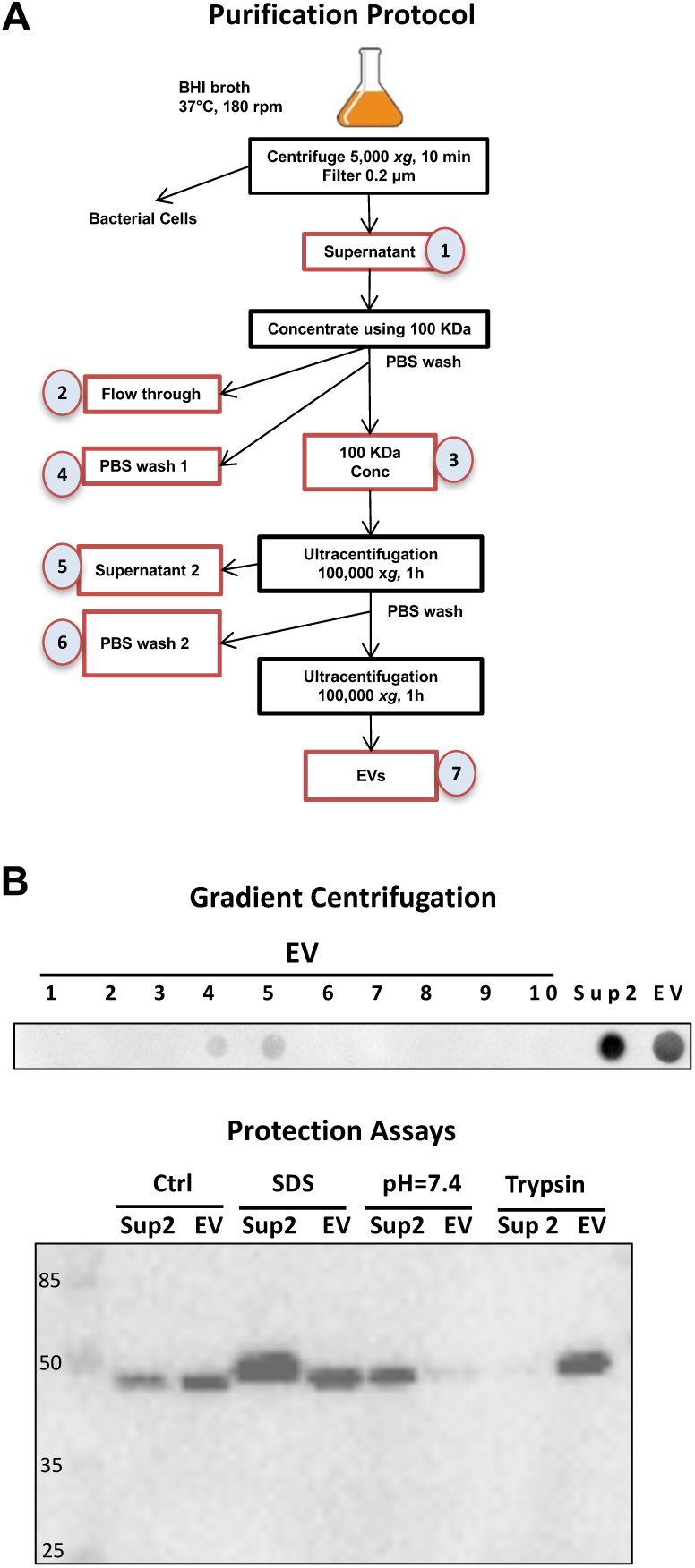
**A**) Diagram illustrating purification of EV. Red boxes are purification steps from where an aliquot was taken. Black boxes are processing steps of the protocol to isolate EVs. (**B**) EVs were subject to gradient centrifugation (top) and protection assays (bottom) to discard the possibility of LLO aggregation and to determine if LLO could be protected by EVs, respectively. Fractions from gradient centrifugations were spotted onto PVDF membrane and probed for LLO. For the Protection Assays the lOOKda Cone (step 3) was subject to the indicated treatment. EVs were extracted. Sup 2 was concentrated using 10 KDa ultrafiltration membranes and blotted for LLO with corresponding Evs.

**Supplemental Figure 2.**
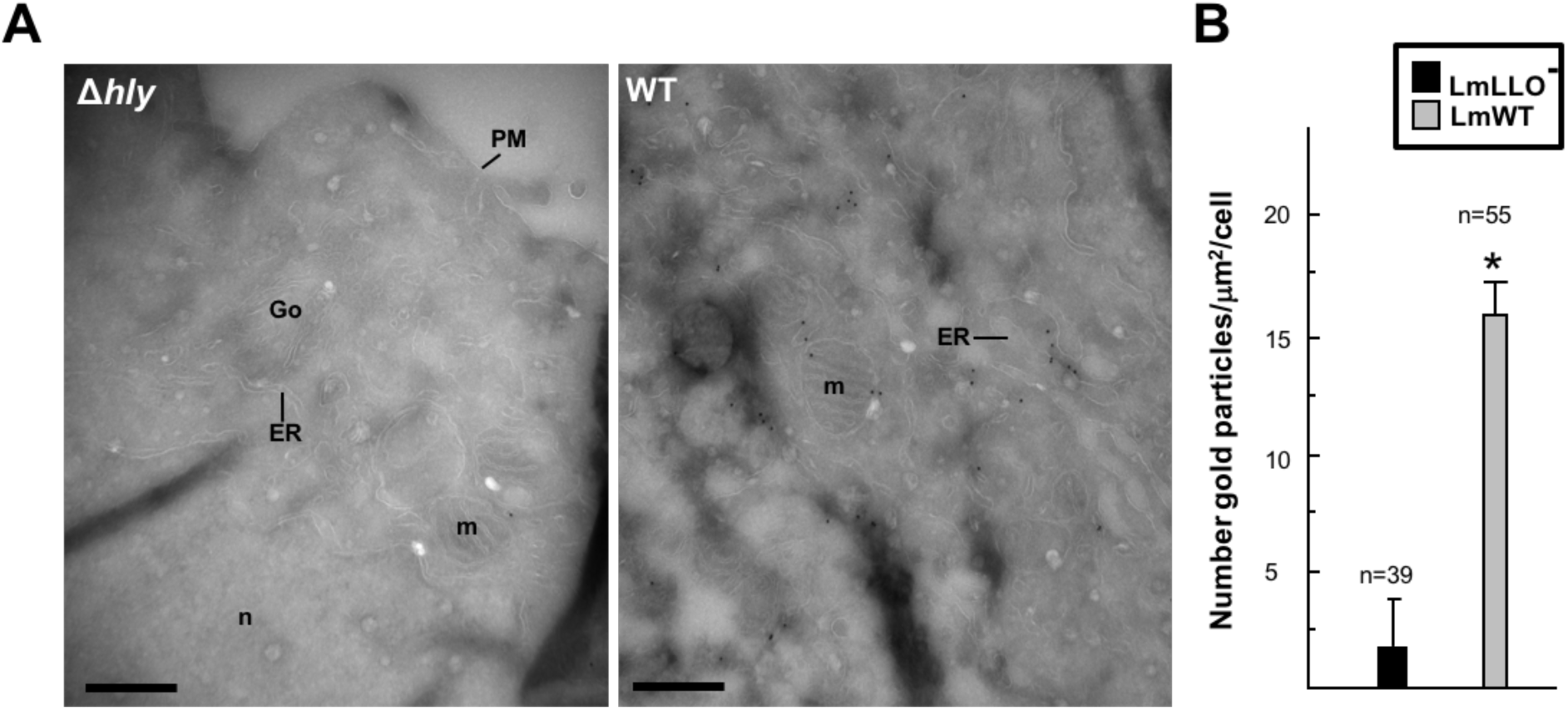
Specificity of the immunostaining for LLO-gold particles. (**A**) Representative views of host cells infected with *L. monocytogenes*-lacking LLO (LmLLO^-^, *Δhly*) or parental strain (LmWT), showing multiple host organelles stained for LLO after infection with wild-type in contrast to infection with LLO^-^. Scale bars 2 μm. (**B**) Gold particle densities (gold particles per μm^2^) per individual cryosections of cells infected with *L. monocytogenes*. Error bars are means ± S.E.M. Mean values differed significantly as determined by the Student’s t-test (n=number of cells analyzed). \**p*<0.0005.

## REFERENCES

1. Lee, E.-Y., Choi, D.-Y., Kim, D.-K., Kim, J.-W., Park, J. O., Kim, S., Kim, S.-H., Desiderio, D. M., Kim, Y.-K., Kim, K.-P., and Gho, Y. S. (2009) Gram-positive bacteria produce membrane vesicles: Proteomics-based characterization of Staphylococcus aureus-derived membrane vesicles. Proteomics. 9, 5425–5436

2. Lecuit, M. (2005) Understanding how Listeria monocytogenes targets and crosses host barriers. Clin. Microbiol. Infect. 11, 430–436

3. Thay, B., Wai, S. N., and Oscarsson, J. (2013) Staphylococcus aureus α-toxin-dependent induction of host cell death by membrane-derived vesicles. PloS one. 8, e54661

4. Olaya-Abril, A., Prados-Rosales, R., McConnell, M. J., Martín-Peña, R., González-Reyes, J. A., Jiménez-Munguía, I., Gómez-Gascón, L., Fernández, J., Luque-García, J. L., García-Lidón, C., Estévez, H., Pachón, J., Obando, I., Casadevall, A., Pirofski, L.-A., and Rodríguez-Ortega, M. J. (2014) Characterization of protective extracellular membrane-derived vesicles produced by Streptococcus pneumoniae. J Proteomics. 106, 46–60

5. Lecuit, M., Ohayon, H., Braun, L., Mengaud, J., and Cossart, P. (1997) Internalin of Listeria monocytogenes with an intact leucine-rich repeat region is sufficient to promote internalization. Infect. Immun. 65, 5309–5319

6. Deatherage, B. L., and Cookson, B. T. (2012) Membrane vesicle release in bacteria, eukaryotes, and archaea: a conserved yet underappreciated aspect of microbial life. Infect. Immun. 80, 1948–1957

7. Camejo, A., Carvalho, F., Reis, O., Leitão, E., Sousa, S., and Cabanes, D. (2011) The arsenal of virulence factors deployed by Listeria monocytogenes to promote its cell infection cycle. Virulence. 2, 379–394

8. Hamon, M. A., Ribet, D., Stavru, F., and Cossart, P. (2012) Listeriolysin O: the Swiss army knife of Listeria. Trends in microbiology. 20, 360–368

9. Vadia, S., Arnett, E., Haghighat, A.-C., Wilson-Kubalek, E. M., Tweten, R. K., and Seveau, S. (2011) The pore-forming toxin listeriolysin O mediates a novel entry pathway of L. monocytogenes into human hepatocytes. PLoS pathogens. 7, e1002356

10. Sawyer, R. T., Drevets, D. A., Campbell, P. A., and Potter, T. A. (1996) Internalin A can mediate phagocytosis of Listeria monocytogenes by mouse macrophage cell lines. Journal of leukocyte biology. 60, 603–610

11. Portnoy, D. A., Jacks, P. S., and Hinrichs, D. J. (1988) Role of hemolysin for the intracellular growth of Listeria monocytogenes. The Journal of experimental medicine. 167, 1459–1471

12. Rivera, J., Cordero, R. J. B., Nakouzi, A. S., Frases, S., Nicola, A., and Casadevall, A. (2010) Bacillus anthracis produces membrane-derived vesicles containing biologically active toxins. Proceedings of the National Academy of Sciences of the United States of America. 107, 19002–19007

13. Poussin, M. A., Leitges, M., and Goldfine, H. (2009) The ability of Listeria monocytogenes PI-PLC to facilitate escape from the macrophage phagosome is dependent on host PKCbeta. Microb. Pathog. 46, 1–5

14. Camilli, A., Goldfine, H., and Portnoy, D. A. (1991) Listeria monocytogenes mutants lacking phosphatidylinositol-specific phospholipase C are avirulent. The Journal of experimental medicine. 173, 751–754

15. Smith, G. A., Marquis, H., Jones, S., Johnston, N. C., Portnoy, D. A., and Goldfine, H. (1995) The two distinct phospholipases C of Listeria monocytogenes have overlapping roles in escape from a vacuole and cell-to-cell spread. Infect. Immun. 63, 4231–4237

16. Gaillard, J. L., Berche, P., Mounier, J., Richard, S., and Sansonetti, P. (1987) In vitro model of penetration and intracellular growth of Listeria monocytogenes in the human enterocyte-like cell line Caco-2. Infect. Immun. 55, 2822–2829

17. Lee, J. H., Choi, C.-W., Lee, T., Kim, S. I., Lee, J.-C., and Shin, J.-H. (2013) Transcription factor σB plays an important role in the production of extracellular membrane-derived vesicles in Listeria monocytogenes. PloS one. 8, e73196

18. Kocks, C., Gouin, E., Tabouret, M., Berche, P., Ohayon, H., and Cossart, P. (1992) L. monocytogenes-induced actin assembly requires the actA gene product, a surface protein. Cell. 68, 521–531

19. Tilney, L. G., and Portnoy, D. A. (1989) Actin filaments and the growth, movement, and spread of the intracellular bacterial parasite, Listeria monocytogenes. The Journal of cell biology. 109, 1597–1608

20. Jiang, Y., Kong, Q., Roland, K. L., and Curtiss, R. (2014) Membrane vesicles of Clostridium perfringens type A strains induce innate and adaptive immunity. Int. J. Med. Microbiol. 304, 431–443

21. Beveridge, T. J. (1999) Structures of gram-negative cell walls and their derived membrane vesicles. Journal of bacteriology. 181, 4725–4733

22. MacDonald, I. A., and Kuehn, M. J. (2012) Offense and defense: microbial membrane vesicles play both ways. Res. Microbiol. 163, 607–618

23. Brown, L., Kessler, A., Cabezas-Sanchez, P., Luque-García, J. L., and Casadevall, A. (2014) Extracellular vesicles produced by the Gram-positive bacterium Bacillus subtilis are disrupted by the lipopeptide surfactin. Molecular microbiology. 93, 183–198

24. Rodrigues, M. L., Nakayasu, E. S., Oliveira, D. L., Nimrichter, L., Nosanchuk, J. D., Almeida, I. C., and Casadevall, A. (2008) Extracellular Vesicles Produced by Cryptococcus neoformans Contain Protein Components Associated with Virulence. Eukaryotic cell. 7, 58–67

25. Brown, L., Wolf, J. M., Prados-Rosales, R., and Casadevall, A. (2015) Through the wall: extracellular vesicles in Gram-positive bacteria, mycobacteria and fungi. Nat Rev Micro. 13, 620–630

26. Kuehn, M. J., and Kesty, N. C. (2005) Bacterial outer membrane vesicles and the host–pathogen interaction. Genes & development

27. Sun, Y., Wilkinson, B. J., Standiford, T. J., Akinbi, H. T., and O’Riordan, M. X. D. (2012) Fatty acids regulate stress resistance and virulence factor production for Listeria monocytogenes. Journal of bacteriology. 194, 5274–5284

28. Mastronicolis, S. K., Arvanitis, N., Karaliota, A., Magiatis, P., Heropoulos, G., Litos, C., Moustaka, H., Tsakirakis, A., Paramera, E., and Papastavrou, P. (2008) Coordinated regulation of cold-induced changes in fatty acids with cardiolipin and phosphatidylglycerol composition among phospholipid species for the food pathogen Listeria monocytogenes. Applied and environmental microbiology. 74, 4543–4549

29. Frydrychowicz, M., Kolecka-Bednarczyk, A., Madejczyk, M., Yasar, S., and Dworacki, G. (2015) Exosomes - structure, biogenesis and biological role in non-small-cell lung cancer. Scand J Immunol. 81, 2–10

30. Marsollier, L., Brodin, P., Jackson, M., Korduláková, J., Tafelmeyer, P., Carbonnelle, E., Aubry, J., Milon, G., Legras, P., André, J.-P. S., Leroy, C., Cottin, J., Guillou, M. L. J., Reysset, G., and Cole, S. T. (2007) Impact of Mycobacterium ulcerans biofilm on transmissibility to ecological niches and Buruli ulcer pathogenesis. PLoS pathogens. 3, e62

31. Vdovikova, S., Luhr, M., Szalai, P., Nygård Skalman, L., Francis, M. K., Lundmark, R., Engedal, N., Johansson, J., and Wai, S. N. (2017) A Novel Role of Listeria monocytogenes Membrane Vesicles in Inhibition of Autophagy and Cell Death. Front Cell Infect Microbiol. 7, 3363

32. Piercey, M. J., Hingston, P. A., and Truelstrup Hansen, L. (2016) Genes involved in Listeria monocytogenes biofilm formation at a simulated food processing plant temperature of 15°C. Int. J. Food Microbiol. 223, 63–74

33. Nakayasu, E. S., Nicora, C. D., Sims, A. C., Burnum-Johnson, K. E., Kim, Y.-M., Kyle, J. E., Matzke, M. M., Shukla, A. K., Chu, R. K., Schepmoes, A. A., Jacobs, J. M., Baric, R. S., Webb-Robertson, B.-J., Smith, R. D., and Metz, T. O. (2016) MPLEx: a Robust and Universal Protocol for Single-Sample Integrative Proteomic, Metabolomic, and Lipidomic Analyses. mSystems. 1, e00043–16

34. Camilli, A., Tilney, L. G., and Portnoy, D. A. (1993) Dual roles of plcA in Listeria monocytogenes pathogenesis. Molecular microbiology. 8, 143–157

35. Carvalho, F., Sousa, S., and Cabanes, D. (2014) How Listeria monocytogenes organizes its surface for virulence. Front Cell Infect Microbiol. 4, 48

36. Fischer, W., and Leopold, K. (1999) Polar lipids of four Listeria species containing L-lysylcardiolipin, a novel lipid structure, and other unique phospholipids. Int. J. Syst. Bacteriol. 49 Pt 2, 653–662

37. Meyer-Morse, N., Robbins, J. R., Rae, C. S., Mochegova, S. N., Swanson, M. S., Zhao, Z., Virgin, H. W., and Portnoy, D. (2010) Listeriolysin O is necessary and sufficient to induce autophagy during Listeria monocytogenes infection. PloS one. 5, e8610

38. Thumser, A. E., and Storch, J. (2007) Characterization of a BODIPY-labeled fluorescent fatty acid analogue. Binding to fatty acid-binding proteins, intracellular localization, and metabolism. Mol. Cell. Biochem. 299, 67–73

39. Johansson, J., Mandin, P., Renzoni, A., Chiaruttini, C., Springer, M., and Cossart, P. (2002) An RNA thermosensor controls expression of virulence genes in Listeria monocytogenes. Cell. 110, 551–561

40. van Bergenhenegouwen, J., Kraneveld, A. D., Rutten, L., Kettelarij, N., Garssen, J., and Vos, A. P. (2014) Extracellular vesicles modulate host-microbe responses by altering TLR2 activity and phagocytosis. PloS one. 9, e89121

41. Schrempf, H., and Merling, P. (2015) Extracellular Streptomyces lividans vesicles: composition, biogenesis and antimicrobial activity. Microb Biotechnol. 8, 644–658

42. Resch, U., Tsatsaronis, J. A., Le Rhun, A., Stübiger, G., Rohde, M., Kasvandik, S., Holzmeister, S., Tinnefeld, P., Wai, S. N., and Charpentier, E. (2016) A Two-Component Regulatory System Impacts Extracellular Membrane-Derived Vesicle Production in Group A Streptococcus. MBio. 10.1128/mBio.00207-16

43. Huang, Q., Gershenson, A., and Roberts, M. F. (2016) Recombinant broad-range phospholipase C from Listeria monocytogenes exhibits optimal activity at acidic pH. Biochim. Biophys. Acta. 1864, 697–705

44. Geoffroy, C., Gaillard, J. L., Alouf, J. E., and Berche, P. (1987) Purification, characterization, and toxicity of the sulfhydryl-activated hemolysin listeriolysin O from Listeria monocytogenes. Infect. Immun. 55, 1641–1646

45. Zemansky, J., Kline, B. C., Woodward, J. J., Leber, J. H., Marquis, H., and Portnoy, D. A. (2009) Development of a mariner-based transposon and identification of Listeria monocytogenes determinants, including the peptidyl-prolyl isomerase PrsA2, that contribute to its hemolytic phenotype. Journal of bacteriology. 191, 3950–3964

46. Burg-Golani, T., Pozniak, Y., Rabinovich, L., Sigal, N., Nir Paz, R., and Herskovits, A. A. (2013) Membrane chaperone SecDF plays a role in the secretion of Listeria monocytogenes major virulence factors. Journal of bacteriology. 195, 5262–5272

47. Bonnemain, C., Raynaud, C., Réglier-Poupet, H., Dubail, I., Frehel, C., Lety, M.-A., Berche, P., and Charbit, A. (2004) Differential roles of multiple signal peptidases in the virulence of Listeria monocytogenes. Molecular microbiology. 51, 1251–1266

48. Gurung, M., Moon, D. C., Choi, C.-W., Lee, J. H., Bae, Y. C., Kim, J., Lee, Y. C., Seol, S. Y., Cho, D. T., Kim, S. I., and Lee, J.-C. (2011) Staphylococcus aureus produces membrane-derived vesicles that induce host cell death. PloS one. 6, e27958

49. Seveau, S., Bierne, H., Giroux, S., Prévost, M.-C., and Cossart, P. (2004) Role of lipid rafts in E-cadherin-and HGF-R/Met-mediated entry of Listeria monocytogenes into host cells. The Journal of cell biology. 166, 743–753

50. Coconnier, M. H., Lorrot, M., Barbat, A., Laboisse, C., and Servin, A. L. (2000) Listeriolysin O-induced stimulation of mucin exocytosis in polarized intestinal mucin-secreting cells: evidence for toxin recognition of membrane-associated lipids and subsequent toxin internalization through caveolae. Cellular microbiology. 487–504

51. Stavru, F., Bouillaud, F., Sartori, A., Ricquier, D., and Cossart, P. (2011) Listeria monocytogenes transiently alters mitochondrial dynamics during infection. Proceedings of the National Academy of Sciences of the United States of America. 108, 3612–3617

52. Villanueva, M. S., Sijts, A. J., and Pamer, E. G. (1995) Listeriolysin is processed efficiently into an MHC class I-associated epitope in Listeria monocytogenes-infected cells. Journal of immunology. 155, 5227–5233

53. Schnupf, P., Zhou, J., Varshavsky, A., and Portnoy, D. A. (2007) Listeriolysin O secreted by Listeria monocytogenes into the host cell cytosol is degraded by the N-end rule pathway. Infect. Immun. 75, 5135–5147

54. Elluri, S., Enow, C., Vdovikova, S., Rompikuntal, P. K., Dongre, M., Carlsson, S., Pal, A., Uhlin, B. E., and Wai, S. N. (2014) Outer membrane vesicles mediate transport of biologically active Vibrio cholerae cytolysin (VCC) from V. cholerae strains. PloS one. 9, e106731

55. Horstman, A. L., and Kuehn, M. J. (2000) Enterotoxigenic Escherichia coli secretes active heat-labile enterotoxin via outer membrane vesicles. J Biol Chem. 275, 12489–12496

56. Kong, M., Bhattacharya, R. N., James, C., and Basu, A. (2005) A statistical approach to estimate the 3D size distribution of spheres from 2D size distributions. Geol Soc America Bull. 117, 244

57. Bannam, T., and Goldfine, H. (1999) Mutagenesis of active-site histidines of Listeria monocytogenes phosphatidylinositol-specific phospholipase C: effects on enzyme activity and biological function. Infect. Immun. 67, 182–186

58. Wolf, J. M., Rivera, J., and Casadevall, A. (2012) Serum albumin disrupts Cryptococcus neoformans and Bacillus anthracis extracellular vesicles. Cellular microbiology. 14, 762–773

59. Tyanova, S., Temu, T., and Cox, J. (2016) The MaxQuant computational platform for mass spectrometry-based shotgun proteomics. Nature Protocols. 11, 2301–2319

60. Polpitiya, A. D., Qian, W.-J., Jaitly, N., Petyuk, V. A., Adkins, J. N., Camp, D. G., Anderson, G. A., and Smith, R. D. (2008) DAnTE: a statistical tool for quantitative analysis of -omics data. Bioinformatics. 24, 1556–1558

61. Kanehisa, M., Sato, Y., and Morishima, K. (2016) BlastKOALA and GhostKOALA: KEGG Tools for Functional Characterization of Genome and Metagenome Sequences. J. Mol. Biol. 428, 726–731

62. Hiller, K., Hangebrauk, J., Jäger, C., Spura, J., Schreiber, K., and Schomburg, D. (2009) MetaboliteDetector: comprehensive analysis tool for targeted and nontargeted GC/MS based metabolome analysis. Anal. Chem. 81, 3429–3439

63. Kind, T., Wohlgemuth, G., Lee, D. Y., Lu, Y., Palazoglu, M., Shahbaz, S., and Fiehn, O. (2009) FiehnLib: mass spectral and retention index libraries for metabolomics based on quadrupole and time-of-flight gas chromatography/mass spectrometry. Anal. Chem. 81, 10038–10048

64. Xia, J., Sinelnikov, I. V., Han, B., and Wishart, D. S. (2015) MetaboAnalyst 3.0–making metabolomics more meaningful. Nucleic Acids Research. 43, W251–7

65. Kyle, J. E., Crowell, K. L., Casey, C. P., Fujimoto, G. M., Kim, S., Dautel, S. E., Smith, R. D., Payne, S. H., and Metz, T. O. (2017) LIQUID: an-open source software for identifying lipids in LC-MS/MS-based lipidomics data. Bioinformatics. 10.1093/bioinformatics/btx046

66. Pluskal, T., Castillo, S., Villar-Briones, A., and Oresic, M. (2010) MZmine 2: modular framework for processing, visualizing, and analyzing mass spectrometry-based molecular profile data. BMC Bioinformatics. 11, 395

67. Bishop, D. K., and Hinrichs, D. J. (1987) Adoptive transfer of immunity to Listeria monocytogenes. The influence of in vitro stimulation on lymphocyte subset requirements. The Journal of Immunology. 139, 2005–2009

68. Jones, S., and Portnoy, D. A. (1994) Characterization of Listeria monocytogenes pathogenesis in a strain expressing perfringolysin O in place of listeriolysin O. Infect. Immun. 62, 5608–5613

69. Muraille, E., Narni-Mancinelli, E., Gounon, P., Bassand, D., Glaichenhaus, N., Lenz, L. L., and Lauvau, G. (2007) Cytosolic expression of SecA2 is a prerequisite for long-term protective immunity. Cellular microbiology. 9, 1445–1454

